# Using attentive gated neural networks to quantify the impact of non-coding variants on transcription factor binding affinity

**DOI:** 10.1101/2021.07.30.454350

**Authors:** Neel Patel, Haimeng Bai, William S. Bush

## Abstract

A large proportion of non-coding variants are present within binding sites of transcription factors(TFs), which play a significant role in gene regulation. Thus, deriving the impact of non-coding variants on TF binding is the first step towards unravelling their regulatory roles within their associated disease traits. Most of the modern algorithms used for this purpose are based on convolutional neural network(CNN) architectures. However, these models are incapable of capturing the positional effect of different sub-sequences within the TF binding sites on the binding affinity. In this paper, we utilize the attentive gated neural network(AGNet) architecture to build a set of TF-AGNet models for predicting *in vivo* TF binding intensities in the GM12878 lymphoblastoid cells. These models have novel layers capable of deriving the impact of relative positions of different DNA sub-sequences, within a binding site, on TF binding affinity, and of extracting the most relevant prediction features. We show that the TF-AGNet models are able to outperform conventional CNNs for predicting continuous values of TF binding affinity. We also train additional TF-AGNet models for 20 TFs using data from 4 other cell-lines to assess the generalizability of their prediction accuracy. Lastly, we show that the TF-AGNet based models more accurately classify non-coding variants that significantly affect TF binding compared to models based on 7 variant annotation tools. This accuracy can be leveraged to derive gene regulatory roles of millions of non-coding variants across the genome to further examine their mechanistic associations with complex disease traits.

## Introduction

About 70% of all the variants associated with complex disease traits are present within the non-coding portion of the genome^1–4^. Functional relationships between such variants and the disease traits, which are otherwise very difficult to establish, can be derived from capturing their gene regulatory context. As non-coding variants are more often than not found within regulatory elements, such as promoters and enhancers, characterizing their gene regulatory roles can lead to mechanistic understanding of the associated disease traits^3–5^. Furthermore, the aforementioned regulatory elements harbor binding sites for transcription factors(TFs), which drive gene expression regulation by significantly influencing the process of transcription.^6^ Thus, TFs and their binding sites (TFBS) provide essential mechanistic markers to compute the influence of non-coding variants on gene expression regulation and ultimately on disease traits.

Several machine learning methods exist to predict the impact of non-coding variants on TF binding affinity or on TFBS disruption. Such predictive algorithms mostly use TF binding preference, in the form of position weight matrices(PWMs), to compare the probability of a TF binding a given DNA sequence containing the alternate allele to that of the one containing the reference allele corresponding to a non-coding variant^7^. These algorithms either use pre-existing sets of PWMs or train models using ChIP-seq data from resourses like the Encyclopedia of DNA Regulatory Elements(ENCODE)^8^ to learn de novo PWMs^9^. Methods such as FIMO(Find Individual Motif Occurrence)^10^, RSAT(Regulatory Sequence Analysis Tools)^11^, Clover^12^ and QBiC-PRED^13^ score DNA sequences containing reference and alternate alleles based on the modifications to the TF motif. However, since these methods must be pre-trained on a reference panel of variants, they cannot be used to annotate novel TFBS altering variants. To overcome this limitation, modern TFBS variant annotation algorithms use deep learning neural networks to generate predictions. Convolutional neural networks(CNNs) are used in DeepSEA^14^, DeepBIND^15^, DANQ^16^ and DeFine^17^ to predict impact of regulatory variants on TF binding. CNNs process input DNA sequences using kernel filters to scan for TF motifs, and to predict the probability of a TF binding to them. Additionally, both DANQ and DeFine include recurrent neural network (RNN) layers, which process the features extracted by the CNNs to capture the long-distance dependencies within the TFBS sequences. While DeepSEA, DeepBIND and DANQ produce binary predictions for TF binding/non-binding, DeFine predicts continuous TF binding intensities. While the former type of output can help deduce the absolute influence of variants on TFBS, more information regarding the changes in TF binding preference, brought about by the variants, can be gleaned by utilizing the latter type of output.

Although the CNN based algorithms have been shown to outperform most other variant annotation tools, they suffer from two main limitations. First, models based on CNNs alone (such as DeepSEA and DeepBIND) only capture the local features from the input TFBS sequences, and ignore the long distance relationships within them. Second, the input layers of these CNN based models only accept one-hot coded matrices of fixed length DNA sequences, which could lead to a sparse representation of these sequences. To overcome these limitations, recent neural network architectures, such as C-LSTM(CNNs with long short-term memory units)^18^ and AGNet(attentive gated neural networks)^19^, have used novel properties such as k-mer vector representations for the input DNA sequences and gated convolutional and recurrent networks(GCNs and GRNs). The C-LSTM architecture uses k-mer vector embeddings normally associated with natural language processing to represent input DNA sequences in the form of continuous multi-dimensional vectors corresponding to their constituent sub-sequences/k-mers^18^. The AGNet architecture further extend this model to include attention layers along with GCNs and GRNs to control the flow of features being passed between the layers and to extract the most informative local and global features from the input sequences^19^. Both of these models have been shown to perform more accurately than conventional CNNs and the CNN-RNN hybrid models for predicting chromatin accessibility based on DNA sequence inputs^18,19^. However, the AGNet model outperforms the C-LSTM in this prediction task due to additional layers that aid in their prediction performance. Despite its success, the AGNet model architecture has not been used for predicting TF binding affinity and annotating the variants that significantly impact it.

In this paper, we have adapted the AGNet architecture to build neural networks, called TF-AGNet, for predicting *in vivo* continuous TF binding intensities utilizing the ENCODE ChIP-Seq datasets corresponding to 149 different TFs(build37) and 144 TFs(build38) for the GM12878 immortalized lymphoblastoid cell line(LCL). We compare the prediction performance of these models to that obtained from the CNN-RNN hybrid models corresponding to the DeFine approach for GM12878 LCL. Additionally, we also train TF-AGNet models for 4 other ENCODE tier-1 and tier-2 cell lines for 20 TFs whose ChIP-Seq data was available across all of these cell lines. Lastly, we assess the accuracy of TF-AGNet models for correctly classifying variants significantly influencing TFBS and compare it to 7 other popular non-coding variant annotation algorithms. Annotating TFBS altering variants, using TF-AGNet models, is the first step towards deriving their gene regulatory functions, which can be used downstream to characterize their mechanistic relationships with complex disease traits.

## Results

### TF-AGNet models were more accurate at predicting TF binding intensity compared to conventional deep learning models

The AGNet model architecture, proposed by Guo et. al.^19^, has been shown to perform better than conventional CNN-RNN hybrid architecture for predicting chromatin accessibility using DNA sequence information in form of k-mer embeddings. It has several novel properties such as :1) Position embedding to capture the relative importance of each k-mer within a TFBS sequence 2) Dual attention layers capable of extracting important information from the sequence and feature based inputs 3) GCN and GRNs for deriving informative local features within and global relatedness among the input k-mers respectively. Here, we have adapted the architecture in order to build TF-AGNet models for predicting *in vivo* ChIP-Seq TF binding intensity values. We trained TF-AGNet models for each one of the 149 TFs(build37/GRCh37 reference assembly) and 144 TFs(build38/GRCh38 reference assembly) using the ChIP-Seq data from the GM12878 immortalized lymphoblastoid cell-line(see **Training the TF-AGNet neural network models** and **Figure 1**). We tested the accuracy of each TF model by calculating Pearson’s Correlation Coefficient (PCC) between observed and predicted scaled log normalized intensity values for a test set of peak regions, which containined 15% of the total regions, selected at random, for each model. The median PCC for all the build37 TF-AGNet models was 0.768, while that for the build38 models was 0.733. Both sets of models had 5 TF models with PCC less than 0.5, out of which 4 low predictive models were common between them (CBX3, CHD4, KDM1A and NFXL1). On the other hand, build37 TF-AGNet models corresponding to 28 TFs were highly accurate (PCC > 0.85), some of which have been shown in **Figure 2A**. For build38, only 12 TF-AGNet models were highly predictive at this threshold of PCC perhaps due to the poorer quality of the ChIP-Seq data aligned to GRCh38 reference build. We have provided the ENCODE accession IDs for all the ChIP-Seq experiments for the two reference builds, along with their evaluation results and the total number of peak regions used to train the models in ***Supplementary table S1***. All the analyses and results presented henceforth will correspond to build37 TF-AGNet models

**Figure 1:**
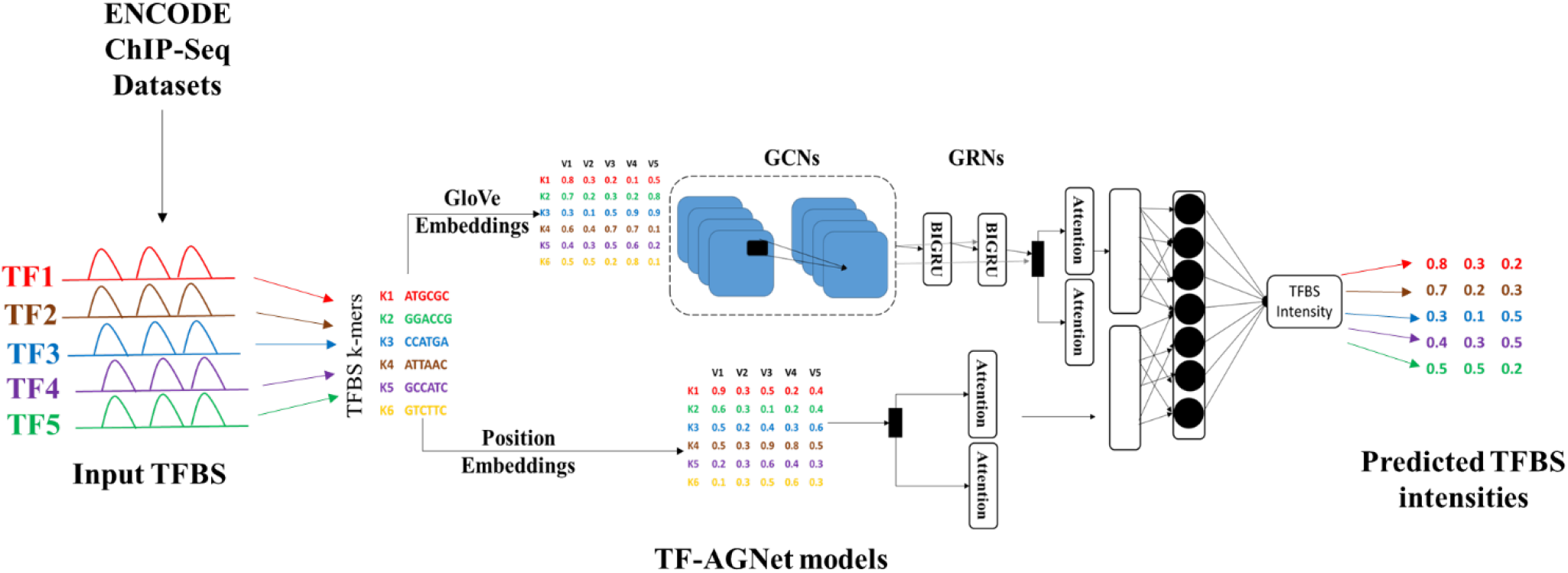
Training the TF-AGNet models for predicting TFBS intensities. We utilized the AGNet architecture to build neural networks for predicting in vivo ChIP-Seq intensities for 149 TFs(GRCh37/build37) and 144 TFs(GRCh38/build38) corresponding to the GM12878 LCL. We downloaded the corresponding processed ChIP-Seq datasets from ENCODE(accession IDs are provided in Supplementary Tables S1A and S1B)

**Figure 2:**
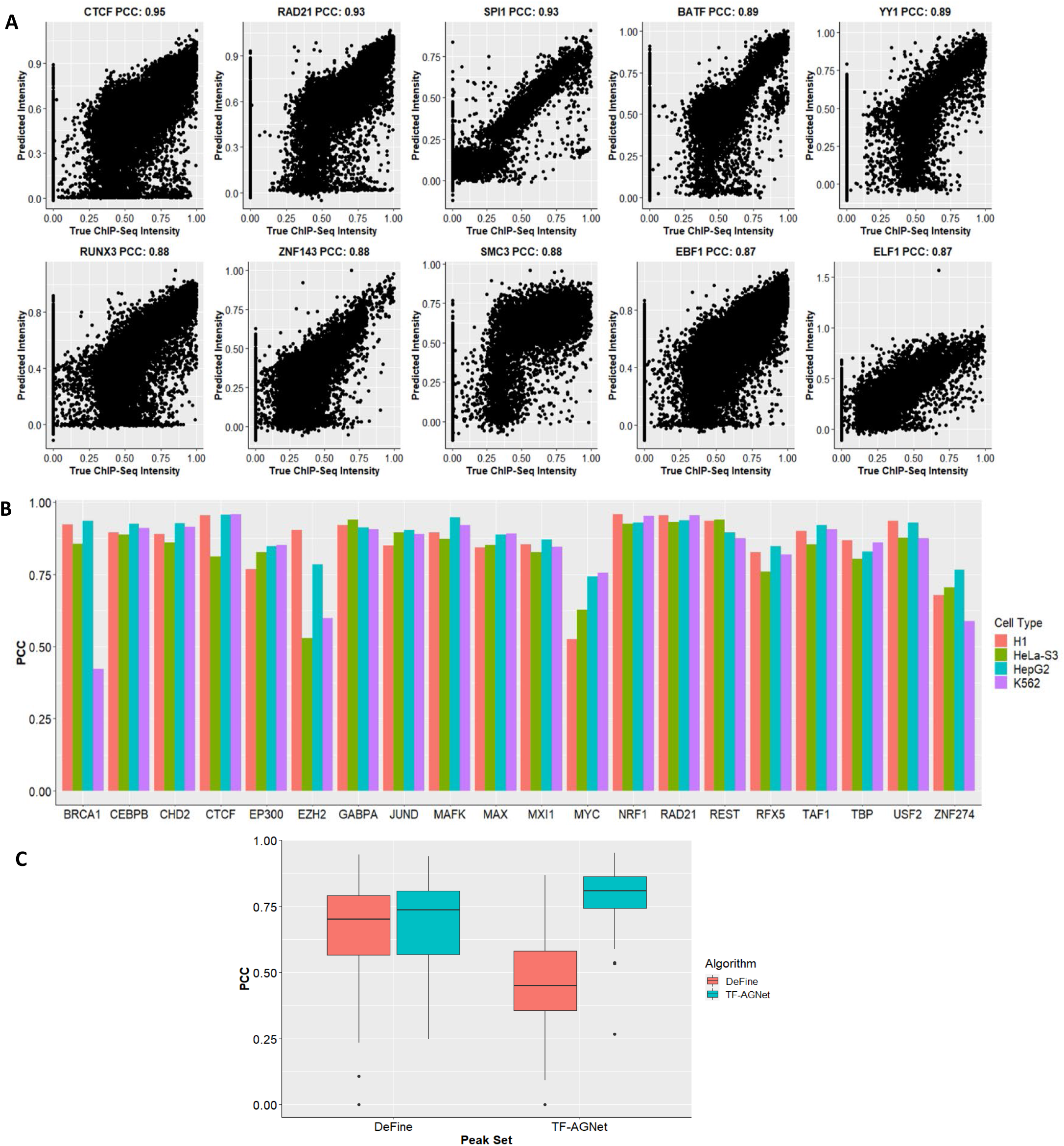
The TF-AGNet models were very accurate at predicting TF binding intensity. A) Scatterplots showing the prediction accuracy f5or the most accurate TF models B) PCC for the 20 TF models trained using the cross-cell type data. C) Boxplots showing the comparison between TF-AGNet and DeFine models from predicting intensity values for TF-AGNet and DeFine peak sets. (****-p-value < 10^−5^)

We also trained models for 4 other cell-lines(K562, H1, HepG2 and HeLa-S3) for 20 TFs, that had data for available for all of these cell-lines in ENCODE using a cross-cell type training strategy described in **Cross-cell type TF-AGNet model training**. As shown in **Figure 2B**, these cross-cell type models were largely accurate(Median-PCC_K562_=0.88, Median-PCC_H1_=0.89, Median-PCC_HepG2_=0.91, Median-PCC_HeLa_=0.85) with the exception being BRCA1-K562 (PCC = 0.422). The detailed evaluation results for these TF-AGNet models have been provided in ***Supplementary table S2***.

We compared the performance of the build37 TF-AGNet models to those built using the DeFine architecture, consisting of a pair of identical conventional CNN layers reading the input DNA sequence for a TFBS in the forward and reverse directions to produce features passed on to a set of fully connected layers for predicting in vivo ChIP-Seq intensity^17^. We hypothesized that the capability of TF-AGNet models to capture positional k-mer information along with increased attention on predictive features would result in more accurate prediction of TF binding intensity compared to that obtained from the DeFine models. In order to test this hypothesis, we predicted intensity values using both models for a set of peak regions used to train DeFine models and a set of regions used to train the TF-AGNet models for 67 GM12878 TFs. As shown in **Figure 2C**, both set of models perform statistically similarly on the DeFine peak regions(Median-PCC_TF-AGNet_ = 0.736; Median-PCC_DeFine_ = 0.701, Wilcoxon p-value = 0.07). However, the TF-AGNet models outperformed the DeFine models significantly(Median-PCC_TF-AGNet_ = 0.806; Median-PCC_DeFine_ = 0.449, Wilcoxon p-value = 6.9e-12), when predictions were made on the peak regions used for training TF-AGNet models. Thus, the TF-AGNet TF models were very accurate at predicting *in vivo* TF binding intensity values, while outperforming the conventional CNN based DeFine models.

### Variants altering TF binding sites *in vivo* were more accurately classified by the TF-AGNet models compared to other variant annotation algorithms

We next assessed the ability of the TF-AGNet models to classify variants that significantly influence binding of TFs leading to an allele specific binding (ASB) event. Such ASB events can correspond to either increase (Gain-of-Binding variants) or a decrease (Loss-of-Binding variants) in the binding affinity. We used previously published *in vivo* differential TF binding changes based on allele specific binding analyses^20^. Furthermore, we compared the performance of the TF-AGNet models to 7 TFBS altering variant annotation tools using data compiled by Wagih et. al.^20^

We identified 10 TFs for which models were available for all the algorithms and used pre-compiled scores (except for the QBiC-Pred method) to classify Gain-of-Binding(GOB) and Loss-of-Binding(LOB) ASB events(see **Determining the accuracy of the TF-AGNet models for predicting allele specific binding(ASB) TF binding events**). We applied three different thresholds(25%, 20%, and 15%) on both ends of the distribution of these scores to call Gain-of-Binding and Loss-of-Binding events for each algorithm. As shown in **Figure 3**, the average AUROC for the TF-AGNet models was the highest among all the 8 algorithms, for the three thresholds for both GOB (AUROC_15_ = 0.647, AUROC_20_ = 0.745, AUROC_25_ = 0.737) and LOB (AUROC_15_ = 0.702, AUROC_20_ = 0.738, AUROC_25_ = 0.732) ASB events. Additionally, while methods such as DeepSEA^14^(LOB-AUROC_20_ = 0.659; GOB-AUROC_20_ =0.674) 0.704),DeepBind^15^(LOB-AUROC_20_ = 0.68; GOB-AUROC_20_ = 0.704), deltaSVM^21^(LOB-AUROC_20_ = 0.702; GOB-AUROC_20_ = 0.714)and QBiC-Pred^13^(LOB-AUROC_20_ = 0.693; GOB-AUROC_20_ = 0.684) were not as accurate as the TF-AGNet models, they still outperformed models based on JASPAR(LOB-AUROC_20_ = 0.634; GOB-AUROC_20_ = 0.630) and MEME(LOB-AUROC_20_ = 0.615; GOB-AUROC_20_ = 0.599) based PWM scores and those based on GERV^22^(LOB-AUROC_20_ = 0.552; GOB-AUROC_20_ = 0.417).

**Figure 3:**
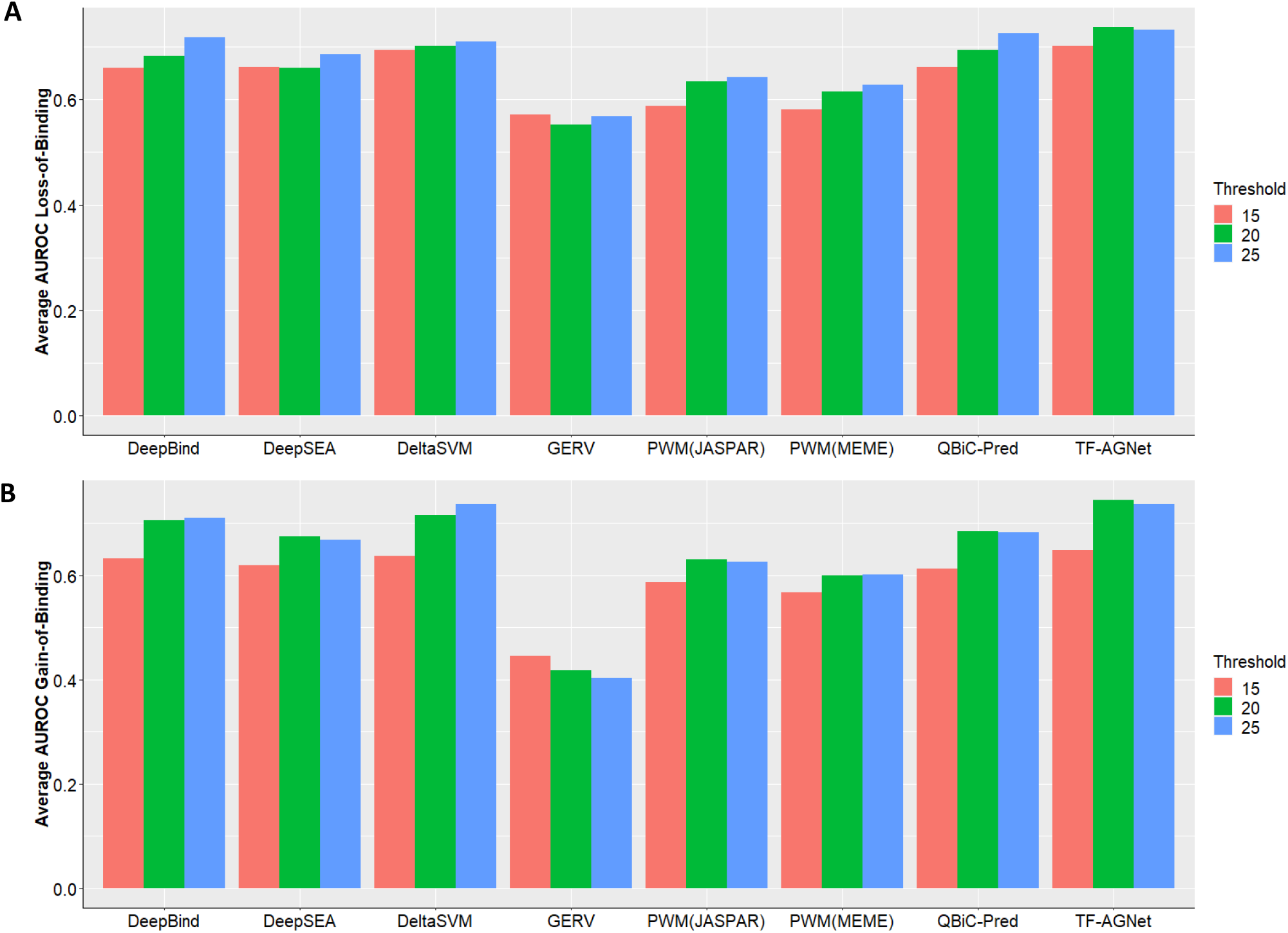
TF-AGNet models were more accurate than other methods for classifying ASB events: Barplots showing the accuracy obtained from classifying ASB events, by the means of average AUROC calculated over 10 different TF models, using different methods for A) 947 Loss-of-Binding events and B) 968 Gain-of-Binding events.

Thus, the TF-AGNet models were superior at predicting ASB specific binding events brought about by the presence of variants within the TFBS in comparison to other variant annotation tools.

## Discussion

In this paper, we trained TF-AGNet models to predict *in vivo* TF binding intensities for the two human genome reference assemblies (build37 and build38) and subsequently used them to quantify the influence of non-coding variants on TF binding in the GM12878 LCL. The observation that the poorly performing TF-AGNet models for the two reference builds corresponded to mostly the same set of TFs, could point towards poor quality of their ChIP-Seq data or could mean that these TFs act as co-factors which don’t directly bind the DNA sequences. Our models were more accurate at predicting TF binding intensity, compared to the CNN-RNN hybrid DeFine models. This improved performance of the TF-AGNet models is mainly due to their capability to capture local and global features from the input k-mers using GCNs and GRNs. The positional k-mer embeddings used in the TF-AGNet models also further add to their improvement by capturing the importance of the k-mers within each TFBS sequence with respect to their relative position. Thus, information beyond motif composition, usually not captured by traditional TFBS prediction models, could also prove to be useful for generating more accurate predictions. Additionally, we observed that the DeFine models, although highly predictive of the intensity values for the TFBS used to originally train them, were not very accurate at predicting intensities for the ones used in this paper. This could potentially be due to overfitting of these models for the specific regions used to train them; a phenomenon not observed for the TF-AGNet models.

We note that TF-AGNet model accuracy comes at a cost of interpretability. Due to the complicated architecture of these models, deriving motif information from them is extremely difficult. CNN models such as DeepSEA, DeepBind and DeFine are better suited for motif discovery analysis due to their simpler and more interpretable architectures. Here, our main focus was to develop accurate TF-variant annotation models, therefore the limited ability to estimate TF binding motifs from TF-AGNet models was not seen as a drawback.

The primary cell-line of focus for training the models was the blood derived GM12878 LCL. However, ENCODE also contains a large number of ChIP-Seq datasets corresponding to other commonly used cell lines. Leveraging this data is extremely useful for building more generalizable prediction models capable of capturing TF binding patterns across different cell-lines/tissues. Furthermore, such generalizable models can be pivotal to examine the functional gene regulatory roles of the non-coding variants in different cell-line/tissue contexts. To facilitate these studies, we have trained models corresponding to 4 other tier-1 and tier-2 ENCODE cell lines for 20 TFs. Our cross-cell type training strategy can be used to generate even more models across different cell lines.

The TF-AGNet models classified variant impact on TF binding better than many commonly used non-coding variant annotation algorithms. The improved performance is likely due to the novel layers used in the AGNet architecture which are capable of capturing the influence of variants over sequential and spatial aspects of TF binding which are not often jointly analyzed by other algorithms. Additionally, we noted that the machine learning-based variant annotation methods significantly outperformed those using PWMs derived from MEME or JASPAR. This was not surprising as the PWMs capture a very limited number of TF binding features^13^.

There are several downstream analyses which could be performed using the highly accurate TF-AGNet based TFBS disruption scores for variants. The most straightforward of which is identifying the pathogenicity of the TFBS altering variants by using the annotation scores to distinguish between variants associated with pathogenic conditions and the neutral variants. Large-scale sequencing studies such as the trans-omics for precision medicine(TopMED)^23^ and the UK-BioBank^24^ have collected whole genome sequences for thousands of individuals. The TF-AGNet models presented in this paper could be used to leverage these rich data sources to derive the functional relationship between millions of non-coding regulatory variants and complex phenotypic traits. To that end, we have built a docker container(https://hub.docker.com/repository/docker/bushlab/tfagnet), which can be used to compute the influence of non-coding variants on the binding sites of the GM12878 TFs based on GRCh37 and GRCh38 TF-AGNet models.

## Methods

### Training the TF-AGNet neural network models

In order to predict the variant effects on TF binding, we trained neural network models, utilizing the attentive gated neural network (AGNet) architecture described by Guo et. al^19^, that predict the TF binding intensity using DNA sequence information. We used processed ChIP-Seq data downloaded from ENCODE, described in ***Supplementary tables S1A*** and ***S1B***, corresponding to the GM12878 LCL in order to train the models as shown in **Figure 1**. We used data corresponding to autosomes (chromosomes 1-22) and aligned to either the build GRCh37/hg19 or build GRCh38/hg38 human genome reference assemblies for training the models. Following steps were involved in pre-processing the data before training the models:

1. For each TF, we first removed the peak regions corresponding to top 1% intensity values, as they represent binding regions with low complexity^17^. We further applied Log10 normalization to the intensity values and scaled them in the range (0,1).
2. Furthermore, we set the maximum length of the peak regions at 2000bp and trimmed the longer regions, on both ends, to bring them to this length.
3. We then downloaded the 1x normalized DNAase-seq tracks for GM12878 trimmed down to the first nucleotide from the 5’ used in the ENCODE dream challenge. Using this data, we filtered out the low accessibility peak regions corresponding to the bottom 10% with respect to DNAase-seq intensity.
4. The above step gave us a positive set of peak regions for training the models pruned by their DNA accessibility. In addition to these, we created a negative set of genomic regions not bound by any TF, which followed the same length and chromosome distribution as that of the positive set for each TF. The ratio of the positive and negative regions was kept at 1.0.
5. Lastly, we downloaded build 37 sequences for each peak region and corrected them for the presence of NA12878 genomic variants using the “FastaAlternateReferenceMaker” tool of GATK(version: 4.1.9) and corresponding VCF file (https://ftp-trace.ncbi.nlm.nih.gov/ReferenceSamples/giab/release/NA12878_HG001/latest/GRCh37/HG001_GRCh37_GIAB_highconf_CG-IllFB-IllGATKHC-Ion-10X-SOLID_CHROM1-X_v.3.3.2_highconf_PGandRTGphasetransfer.vcf.gz). This corrected for the NA12878 specific variants present within each peak region.
6. The set of sequences generated above were then used to train the GloVe model^25^ in order to learn vector embeddings for each sub-sequence/k-mer. In order to accomplish this, we divided each TFBS sequence into k-mers of length 6 and stride 2, which were then aggregated into a big corpus of k-mers containing sequence information regarding all the TFBS for 149 TFs. This corpus contained 4,258 unique k-mers. We trained the GloVe model using the github code(https://github.com/stanfordnlp/GloVe) using the vector size of 100 for 100 iterations`.

After generating the GloVe vector embeddings for the k-mers, we utilized the AGNet architecture with the keras package(v 2.3.1) and the tensorflow (v 1.14.0) backend to train the neural network models. We trained individual models for each TF using the k-mer vector embeddings corresponding to its peak regions as input and the normalized scaled ChIP-Seq intensities as the output. Below we have described the architecture of TF-AGNet briefly, and we refer the readers to the AGNet paper^19^ for further details.

The first layer of the TF-AGNet models was an embedding layer receiving inputs in form of indexed k-mer vectors for each TF peak region. The embedding layer was followed by a layer of multi-scale CNN layer of three 1D CNNs all connected to the embedding layer and learning local informative features from the input k-mers in parallel. Each one of the convolutional layers contained 64 filters and the kernel size for them was 3,5 and 7. Each multi-scale CNN was followed by a max-pooling layer with a pooling size of 3.

Apart from the multi-scale CNNs, the embedding layer was also connected to a position embedding layer that contained vector embeddings for the k-mers derived based on their positions within each input sequence. The position embeddings were of the similar size(100) to that of the GloVe based k-mer embeddings and the two were added to produce the output of the position embedding layer. The output of the position embedding is then passed on to a dual attention layer meant to extract important sequence features.

The outputs from the multi-scale CNNs were concatenated and were passed on to gated convolutional network (GCN) layer. The GCN layer consisted of a conventional 1D CNN with a “sigmoid” activation function and a novel 1D CNN with a scaled exponential linear unit(SELU) activation function for improved gating and control of information flow in form of features to the next layer. Both the CNNs in the GCN layer contained 192 filters and the kernel size of 3. The outputs from these two CNNs were multiplied and a maxpooling of size 3 was applied to product.

The GCNs were followed by two gated recurrent network(GRN) layers containing stacked bi-directional gated recurrent units(BIGRU). Each GRN layer contained 256 nodes and the output dimension was 128. The output from the GCN layer is directly passed on to both the GRNs, and it is also concatenated to their outputs.

The GRN-GCN concatenated output is passed onto another dual attention layer to extract important abstract features. These features are then concatenated to the ones obtained from the dual attention layer containing position embeddings. The concatenated sequence and abstract features are then passed on to a fully connected dense layer containing 128 nodes and the SELU activation function, which is then connected to an output node without any activation function to produce a continuous output corresponding to the TFBS intensity.

The parameters for the TF-AGNet models are initialized using the Lecun normal initializer. Each TF-AGNet model was trained using the mean squared error(MSE) loss and optimized using the Adam optimizer for a maximum of 100 epochs. Furthermore, overfitting in each model was controlled using dropout layers, L2 regularization and early stopping after seeing no improvement in validation loss for 10 straight epochs.

Each TF peak region set containing negative and positive sets was divided into 70%-15%-15% training, validation and test set. After model training was finished, the TF binding intensities of the test set were predicted and were correlated with the actual binding intensities to evaluate the accuracy of the models.

### Cross-cell type TF-AGNet model training

In order to generate generalizable TF-AGNet models, which could be applied to TF binding data corresponding to multiple different cell-types, we trained them using a cross-cell type training strategy. Specifically, we downloaded ChIP-Seq data from the Tier-1 cell lines (K562 immortalized chronic myelogenous leukemia cell line and H1 human embryonic stem cell line) and Tier-2 cell lines(HeLa-S3 immortalized cervical cancer cell line and HepG2 liver carcinoma cell line). We identified 20 TFs, whose data was available for these 4 cell lines and processed it using the steps described in **Training the TF-AGNet neural network models**. We then used the processed data for cross-cell type training using the following procedure: We divided the total processed data for each TF for each one of the 4 cell lines into 70-15-15 training-test-validation set. We then pooled the validation set for all the 4 cell-lines together. We began training the models for each cell-line for each TF using the corresponding pre-trained GM12878 TF-AGNet model. We used training data specific to each cell-line and the validation data from the other three cell-lines to train each TF model. Thus, we trained 4 different models, corresponding to the 4 cell-lines, for each TF, where the training data was cell-line specific and the validation data was obtained from the other 3 cell lines. The training parameters for the cross-cell type training were the same as that used for the GM12878 model training in **Training the TF-AGNet neural network models**. We assessed the accuracy for each TF cell line model using the test data specific to that cell line.

### Comparison of TF-AGNet models with the DeFine models

DeFine models are a set of CNNs trained using the ENCODE ChIP-Seq datasets to predict TF binding intensity^17^. We compared the accuracy of our TF-AGNet models for predicting TF binding intensity to that obtained from the DeFine models. We first downloaded DeFine models corresponding to 67 TFs, for which trained TF-AGNet models were available in our dataset, for the GM12878 cell line. We also downloaded the peak sequences and normalized TF binding intensity used to train the DeFine models. We then selected 15% of the peak regions used to train the DeFine models and predicted their intensity values using both DeFine and TF-AGNet models. We did the same thing with the peak regions used to train the TF-AGNet models. We compared the prediction accuracy of the two types of models for each TF by calculating the PCC between the observed and the predicted intensities for the two different sets of peak regions.

### Determining the accuracy of the TF-AGNet models for predicting allele specific binding(ASB) TF binding events

In order to assess the accuracy of the TF-AGNet models for classifying allele specific(ASB) TF binding events, we downloaded the ASB data aggregated by Wagih et al.^20^ based on differential TF binding identified by ChiP-Seq experiments for 81 TFs. In this dataset, there were 32,252 ASB events (P_binomial_ < 0.01) and 79,827 non-ASB events(P_binomial_ > 0.5). We compared the performance of TF-AGNet models to 7 other methods: QBiC-Pred^13^, DeepSEA^14^, DeepBind^15^, PWM(Meme)^10^, PWM(Jaspar)^26^, GERV^22^ and deltaSVM^21^. We identified 1,915 ASB events(968 Gain-of-Binding and 947 Loss-of-Binding) corresponding to 613 single nucleotide polymorphisms(SNPs) variants and 10 TFs for which prediction models were present for all the algorithms. For these 10 TFs, we had 2,170 non-ASB events(1132 Gain-of-Binding and 1038 Loss-of-Binding) corresponding to 1,001 SNPs. We scored both the ASB and non-ASB SNPs using the TF-AGNet models by first centering the variants within the TFBS identified for the 10 TFs in our original GM12878 based peak set. We then generate a pair of sequences for each variant containing an alternate allele(S_ALT_) and a reference allele(S_REF_) and scored both the sequences. The variant influence on the TFBS(S_v_) was then derived from the difference in the two scores as shown in equation (1).

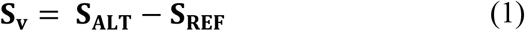

We also scored the variants using the QBiC-Pred algorithm and used the z-scores as the ASB and non-ASB variant influence on TF binding. For the remaining 6 methods, we simply used the scores compiled by Wagih et. al.^20^ for each ASB and non-ASB variant for the 10 TFs. Furthermore, for each algorithm, we used thresholds of 25^th^, 20^th^ and 15^th^ percentiles in order to classify the ASB events based on the scores such that events with scores in the top portion of that percentile were considered Gain-of-Binding ASB events, while the events in the bottom portion of the percentile were considered Loss-of-Binding ASB events. The remaining events were considered as non-ASB events. We then used AUROC to calculate the accuracy of each algorithm for correctly identifying an ASB event using the ground truths from the data compiled by Wagih et. al.^20^

## References

1. Buniello A, MacArthur JAL, Cerezo M, et al. The NHGRI-EBI GWAS Catalog of published genome-wide association studies, targeted arrays and summary statistics 2019. Nucleic Acids Res. 2019;47(D1):D1005–D1012. doi:10.1093/nar/gky1120

2. Claussnitzer M, Cho JH, Collins R, et al. A brief history of human disease genetics. Nature. 2020;577(7789):179–189. doi:10.1038/s41586-019-1879-7

3. Bryzgalov LO, Antontseva E V, Matveeva MY, et al. Detection of regulatory SNPs in human genome using ChIP-seq ENCODE data. PLoS One. 2013;8(10):e78833. doi:10.1371/journal.pone.0078833

4. Farh KK-H, Marson A, Zhu J, et al. Genetic and epigenetic fine mapping of causal autoimmune disease variants. Nature. 2015;518(7539):337–343. doi:10.1038/nature13835

5. Maurano MT, Humbert R, Rynes E, et al. Systematic localization of common disease-associated variation in regulatory DNA. Science. 2012;337(6099):1190–1195. doi:10.1126/science.1222794

6. Lambert SA, Jolma A, Campitelli LF, et al. The Human Transcription Factors. Cell. 2018;172(4):650–665. doi:https://doi.org/10.1016/j.cell.2018.01.029

7. Carrasco Pro S, Bulekova K, Gregor B, Labadorf A, Fuxman Bass JI. Prediction of genome-wide effects of single nucleotide variants on transcription factor binding. Sci Rep. 2020;10(1):17632. doi:10.1038/s41598-020-74793-4

8. Davis CA, Hitz BC, Sloan CA, et al. The Encyclopedia of DNA elements (ENCODE): data portal update. Nucleic Acids Res. 2017;46(D1):D794–D801. doi:10.1093/nar/gkx1081

9. Schacht T, Oswald M, Eils R, Eichmüller SB, König R. Estimating the activity of transcription factors by the effect on their target genes. Bioinformatics. 2014;30(17):i401–i407. doi:10.1093/bioinformatics/btu446

10. Grant CE, Bailey TL, Noble WS. FIMO: scanning for occurrences of a given motif. Bioinformatics. 2011;27(7):1017–1018. doi:10.1093/bioinformatics/btr064

11. Thomas-Chollier M, Defrance M, Medina-Rivera A, et al. RSAT 2011: regulatory sequence analysis tools. Nucleic Acids Res. 2011;39(Suppl_2):W86–W91. doi:10.1093/nar/gkr377

12. Frith MC, Fu Y, Yu L, Chen J-F, Hansen U, Weng Z. Detection of functional DNA motifs via statistical over-representation. Nucleic Acids Res. 2004;32(4):1372–1381. doi:10.1093/nar/gkh299

13. Martin V, Zhao J, Afek A, Mielko Z, Gordân R. QBiC-Pred: quantitative predictions of transcription factor binding changes due to sequence variants. Nucleic Acids Res. 2019;47(W1):W127–W135. doi:10.1093/nar/gkz363

14. Zhou J, Troyanskaya OG. Predicting effects of noncoding variants with deep learning– based sequence model. Nat Methods. 2015;12(10):931–934. doi:10.1038/nmeth.3547

15. Alipanahi B, Delong A, Weirauch MT, Frey BJ. Predicting the sequence specificities of DNA-and RNA-binding proteins by deep learning. Nat Biotechnol. 2015;33(8):831–838. doi:10.1038/nbt.3300

16. Quang D, Xie X. DanQ: a hybrid convolutional and recurrent deep neural network for quantifying the function of DNA sequences. Nucleic Acids Res. 2016;44(11):e107. doi:10.1093/nar/gkw226

17. Wang M, Tai C, E W, Wei L. DeFine: deep convolutional neural networks accurately quantify intensities of transcription factor-DNA binding and facilitate evaluation of functional non-coding variants. Nucleic Acids Res. 2018;46(11):e69–e69. doi:10.1093/nar/gky215

18. Min X, Zeng W, Chen N, Chen T, Jiang R. Chromatin accessibility prediction via convolutional long short-term memory networks with k-mer embedding. Bioinformatics. 2017;33(14):i92–i101. doi:10.1093/bioinformatics/btx234

19. Guo Y, Zhou D, Li W, Nie R, Hou R, Zhou C. Attentive gated neural networks for identifying chromatin accessibility. Neural Comput Appl. 2020;32(19):15557–15571. doi:10.1007/s00521-020-04879-7

20. Wagih O, Merico D, Delong A, Frey BJ. Allele-specific transcription factor binding as a benchmark for assessing variant impact predictors. bioRxiv. January 2018:253427. doi:10.1101/253427

21. Lee D, Gorkin DU, Baker M, et al. A method to predict the impact of regulatory variants from DNA sequence. Nat Genet. 2015;47(8):955–961. doi:10.1038/ng.3331

22. Zeng H, Hashimoto T, Kang DD, Gifford DK. GERV: a statistical method for generative evaluation of regulatory variants for transcription factor binding. Bioinformatics. 2016;32(4):490–496. doi:10.1093/bioinformatics/btv565

23. Taliun D, Harris DN, Kessler MD, et al. Sequencing of 53,831 diverse genomes from the NHLBI TOPMed Program. Nature. 2021;590(7845):290–299. doi:10.1038/s41586-021-03205-y

24. Bycroft C, Freeman C, Petkova D, et al. The UK Biobank resource with deep phenotyping and genomic data. Nature. 2018;562(7726):203–209. doi:10.1038/s41586-018-0579-z

25. Pennington J, Socher R, Manning CD. GloVe: Global Vectors for Word Representation. In: Empirical Methods in Natural Language Processing (EMNLP). ; 2014:1532–1543. http://www.aclweb.org/anthology/D14-1162.

26. Fornes O, Castro-Mondragon JA, Khan A, et al. JASPAR 2020: update of the open-access database of transcription factor binding profiles. Nucleic Acids Res. 2019;48(D1):D87–D92. doi:10.1093/nar/gkz1001

